# RNA allelic frequencies of somatic mutations encode substantial functional information in cancers

**DOI:** 10.1101/2023.03.09.531725

**Authors:** James R.M. Black, Thomas P. Jones, Carlos Martínez-Ruiz, Maria Litovchenko, Clare Puttick, Nicholas McGranahan

**Affiliations:** Cancer Research UK Lung Cancer Centre of Excellence, University College London Cancer Institute, London, UK; Cancer Genome Evolution Research Group, Cancer Research UK Lung Cancer Centre of Excellence, University College London Cancer Institute, London, UK; Cancer Evolution and Genome Instability Laboratory, The Francis Crick Institute, London, UK

## Abstract

A central goal of cancer research is the identification of cancer genes that drive tumour growth and progression. Existing approaches to this problem typically leverage frequentist approaches based on patterns of somatic mutagenesis in DNA. Here, we interrogate RNA variant allele frequencies to identify putative cancer genes with a novel computational tool, *RVdriver*, from bulk genomic-transcriptomic data within 7,948 paired exomes and transcriptomes across 30 cancer types. An elevated RNA VAF reflects a signal from multiple biological features: clonal mutations; mutations retained or gained during somatic copy-number alterations; mutations favoured by allele-specific expression; and mutations in genes expressed preferentially by the tumour compartment of admixed bulk samples. *RVdriver*, a statistical approach that classifies RNA VAFs of nonsynonymous mutations relative to a synonymous mutation background, leverages this information to identify known, as well as putatively novel, cancer genes, with comparable performance to DNA-based approaches. Furthermore, we demonstrate RNA VAFs of individual mutations are able to distinguish ‘driver’ from ‘passenger’ mutations within established cancer genes. Low-RNA VAF *EGFR* mutations otherwise annotated as drivers of glioblastoma by DNA tools harbour a phenotype of reduced EGFR signalling, whilst high-RNA VAF *KDM6A* mutations otherwise annotated as passengers exhibit a driver-like H3K27me3 expression profile, demonstrating the value of our approach in phenotyping tumours. Overall, our study showcases a novel approach for cancer gene discovery, and highlights the potential value of multi-omic and systems-biology approaches in finding novel therapeutic vulnerabilities in cancer to bring about patient benefit.

## Introduction

Tumours evolve in a Darwinian fashion, whereby cancer cells acquire heritable variation that acts as the substrate for selection in the context of its environment. Cancer cell phenotypes are profoundly different from those of cells found in non-cancer tissues, and different cancers often share common phenotypes, or hallmarks^1^. According to the Darwinian model of tumour evolution, this divergence from normal cell phenotype occurs as selection acts upon sequentially acquired, heritable variation^2^. The best studied form of such variation is genetic point mutations.

A central goal of cancer research has been to identify which somatic alterations are the key drivers of cancer initiation, development and metastasis. As such, multiple algorithms and computational tools have been developed to identify patterns of excessive mutagenesis and thus cancer genes, which have been causally implicated in cancer. These include approaches to define genes that are more frequently mutated than would be expected by chance^3,4^, or harbour more nonsynonymous mutations relative to an expected background derived by considering the observed mutational processes in a given set of cancer genomes^5–7^. Other algorithms look for genes in which mutations tend to cluster within genes^8–11^, recur in loci encoding specific regions of the 3D protein confirmation^12^, or occur in domains which might be expected to be functionally relevant to the encoded protein^13,14^.

Such efforts are relatively advanced, and a decade of gradual improvement has facilitated the development of machine-learning approaches that leverage genomic output and deliver interpretable assessments of cancer genes and driver mutations^15^.

However, it is notable that the vast majority of tools and approaches to identify cancer genes and driver mutations focus almost exclusively on genomic data. Many genomic studies also collect transcriptomic data^16^. Usage of this data to uncover novel cancer genes is relatively nascent^17^. Some studies utilise paired or unpaired gene expression data, either to alleviate the burden of testing for repeated measures by pruning lists of putative cancer genes to those that are expressed only^5^, or by leveraging the observation that cancer genes tend to be more uniformly expressed than non-cancer genes^18,19^. However, approaches that incorporate transcriptomic information as a bridge between the frequentist approaches to cancer gene discovery that focus exclusively on the genome, and tumour phenotype, are lacking.

Here, we wanted to understand if by incorporating RNA information we could improve our power to identify the drivers of cancer development and shed light on their key features. Using 7,948 paired whole exome-RNA seq samples taken from tumours across 30 cancer types from the Cancer Genome Atlas (TCGA), we set out to understand the capacity of transcriptomic data to contribute to cancer gene identification. We present ‘RNA VAF driver’ (*RVdriver*), an end-to-end cohort-level bioinformatic algorithm to identify cancer genes, which exploits paired WES/WGS and RNA-seq data. Leveraging *RVdriver*, we discovered that the variant allele frequencies of somatic mutations, when observed in RNA, were capable of distinguishing cancer genes from non-cancer genes across multiple cancer types with comparable sensitivity and specificity to a suite of best-in-class tools which leverage DNA information. Furthermore, within individual mutations, signals from RNA VAF provided meaningful information to enhance existing ‘driver’ mutation discovery tools.

## Results

### Mutation expression in solid tumours

We collated a pan-cancer cohort of tumours with paired whole-exome and RNA sequencing from the Cancer Genome Atlas (TCGA). Hypermutator tumours and samples which had a significant degree of RNA degradation were removed^20^. In total, the cohort comprised 7,948 samples across 30 cancer types (Figure S1). Mutations from these samples were derived from the publically available ‘MC3’ mutation table^21^. In total, these tumours collectively harboured 962,580 somatic mutations, which were further filtered to remove single-nucleotide variants that contained within the ENCODE blacklist^22^. Existing usage of transcriptomic data in tools to identify cancer genes has typically focussed on information relating to gene expression amplitude, or inter-sample heterogeneity. Instead, we processed samples to obtain information relating to the expression of individual mutations.

In the first instance, this facilitated analysis of patterns of mutation expression in solid tumours. Overall, 39% of somatic point mutations were expressed (defined as >1 non-duplicated read aligning to the mutated allele at the mutated position) across all samples, whilst at 41% of mutated positions, there was no expression from either the wild-type or mutated allele (defined as < 2 non-duplicated reads aligning to the mutated or wild-type allele at the mutated position). The remaining 20% corresponds to mutated positions at which only the wild-type allele is expressed (Figure S2A). This relationship differed across synonymous, missense and nonsense mutations (Figures 1A, S2B); as well as across genes within or outside of the COSMIC Cancer Gene Census (CGC) list of cancer genes^20,23^ (mutations within these genes might be more likely to be functionally impactful), and was broadly consistent across cancer types (Figures S3, S4).

**Figure 1.**
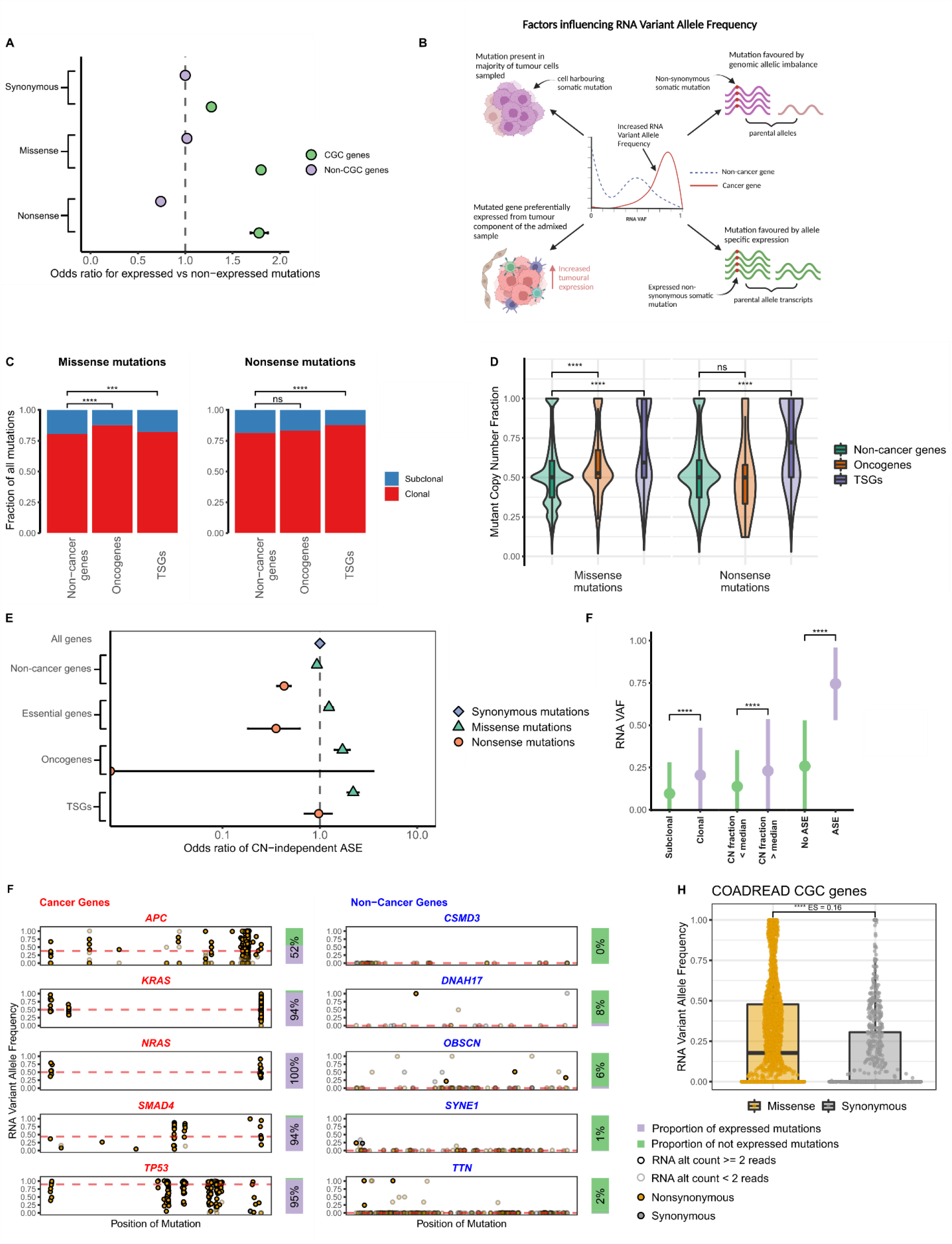
RNA VAF across cancer and non-cancer genes. **A**. Odds ratios for the expression of mutations in genes found, or not found, within the COSMIC cancer gene census (CGC) list. Mutations were identified within the ‘MC3’ mutation table, within samples containing paired RNA sequencing information. For a mutation to be classified as expressed, two or more molecules mapping to the mutated allele had to be identified within RNA. Odds ratios were estimated using the number of expressed and non-expressed mutations observed of a given type, relative to synonymous mutations in non-CGC genes using a Fisher’s exact test; bars indicate 95% confidence intervals for the odds ratio estimate. **B**. Mechanisms of elevated variant allele frequency observed in RNA. **C**. Proportion of all mutations with Cancer Cell Fraction of 1 (i.e. clonal mutations). Fraction of all expressed mutations within the MC3 mutation table which were estimated to be present in 100% of tumour cells within the admixed bulk sample; split by mutation type, and by the function of the gene containing the mutation. **D**. Ratio of the estimated mutated allele copy-number relative to the total copy-number at that position; split by mutation type, and by the function of the gene containing the mutation. **E**. Odds ratio of missense, nonsense and splice site mutations exhibiting allele-specific expression, beyond that expected given the copy-number status of the mutation, and assuming 100% expression from the tumour compartment of the admixed sample. Odds ratio and 95% confidence intervals defined relative to synonymous mutations among all genes. **F**. RNA VAFs of somatic mutations across conditions in C,D and E. Points represent mean RNA VAFs for each condition with lines representing the standard deviation around the mean. **** indicates p<0.0001; Wilcoxon test. **G**. RNA VAFs of somatic mutations within putative cancer genes in colorectal tumours, and RNA VAFs of somatic mutations within putative non-cancer genes in colorectal tumours. Barplot percentages indicate the proportion of nonsynonymous mutations that are expressed. **H**. RNA VAF of synonymous versus non-synonymous mutations within COSMIC cancer gene census genes; **** indicates p<0.0001; Wilcoxon test.

In non-CGC genes, nonsense mutations were less likely to be expressed than synonymous or missense mutations, in keeping with the degradation of these mutated transcripts through nonsense-mediated decay (Figure 1A). Synonymous or missense mutations in these genes were expressed to a similar extent. Of note, though, within CGC genes synonymous mutations were less likely to be expressed than missense mutations, or nonsense mutations within these genes (Figure 1A). This might relate to concurrent copy-number loss of wild-type alleles in the context of deleterious mutations in tumour suppressor genes, where paradoxical overexpression of mutated alleles has previously been described^24^. Regardless of the mechanism, within these genes the likely degradation of truncated transcripts through nonsense-mediated decay does not prevent functionally impactful nonsynonymous mutations, independent of whether they encode a simple base change or a frameshift, from being more frequently expressed than non-impactful mutations. This analysis highlights that mutation expression data can encode information relating to functional relevance of genes.

### RNA VAF

Mutation expression can be considered in conjunction with expression of the wild-type allele. This has utility because in most biological settings, expression of one parental allele positively correlates with expression of the other parental allele^25^. It follows that if functionally important information is encoded in deviation from this pattern, mutated allelic expression changes might be best observed in the context of wild-type allelic expression.

To that end, we leveraged *RVdriver* to calculate RNA variant allele frequencies (RNA VAF) for each mutation, by dividing the number of non-duplicated reads mapping to the variant allele by the total number of non-duplicated reads mapping to either parental allele at that position.

The RNA VAF of each somatic mutation might plausibly be influenced by a variety of factors, some of which relate to the specific mutation, and others that are gene-level or global (Figure 1B). Mutation-level factors that would influence RNA VAF include: the proportion of tumour cells containing the somatic mutation (cancer cell fraction, CCF); the degree of allelic imbalance in genomic copy number state between mutated and non-mutated alleles; and allele-specific expression (where mutated and non-mutated alleles are differentially expressed relative to their genomic copy-number). Gene or sample-level factors include the fraction of the admixed bulk sample derived from the tumour (purity); the degree to which the mutated gene is expressed; and global differences in RNA content between the tumour and non-tumour components of the admixed sample, adjusted for differences in total genomic content^18,26,27^. Thus, a multitude of information might be encoded within a simple, continuous metric, RNA VAF, which to-date has not been used to identify cancer genes.

### RNA VAF differentiates between mutations in cancer genes and non-cancer genes

We next evaluated the extent to which these mutation-level variables influencing RNA VAF differed between mutations in cancer genes and non-cancer genes.

First, we compared the clonal status of mutations between nonsynonymous mutations within oncogenes, tumour suppressor genes (both defined by Bailey et al^20^), and non-cancer genes (genes not found within CGC list or Bailey list). This information has previously been shown to be informative when identifying cancer genes^28^. The clonal status of mutations was calculated using a previously published approach^29^. A large portion of mutations were estimated to have a CCF of 1 (i.e. estimated to be present within 100% of tumour cells within the admixed bulk sample). Missense mutations within oncogenes and tumour suppressor genes were more often clonal than those within non-cancer genes (Figure 1C), whilst nonsense mutations were more likely to be clonal within tumour suppressor genes than non-cancer genes.

Second, we evaluated whether allelic imbalance tended to favour the mutated allele (i.e. mutated alleles were preferentially gained or preserved in copy number events relative to the wild-type allele) more in cancer genes than non-cancer genes. This has been demonstrated to be the case previously in oncogenes^30^, and wild-type copies of tumour suppressor genes are lost canonically in carcinogenesis^31–33^. For each mutation, we calculated its copy-number, estimated using its variant allele frequency in DNA as well as purity and the total copy-number at that position (calculated using ASCAT^34^; for equation see Methods). We then divided this by the total copy-number at the position to obtain a mutant copy-number fraction. Missense mutations within oncogenes, and missense and nonsense mutations within tumour suppressors were typically more likely to be favoured in allelic imbalance, relative to mutations within non-cancer genes (Figure 1D).

Third, we investigated the extent of copy-number independent allele-specific expression (ASE) in mutations within cancer genes relative to non-cancer genes. The observation that expression relative to genomic content is imbalanced between the tumour and non-tumour components of the admixed sample (made in multiple recent papers^18,26,27^) could plausibly confound a direct measurement of allelic expression relative to DNA allele frequencies. We therefore adopted a conservative approach to detect ASE, sensitive to differences between tumour purity and tumour transcript fraction. We considered ASE within non-CGC genes, cancer-cell essential genes, oncogenes and tumour suppressor genes. When measured this way, as expected, ASE favouring the expression of the mutated allele was seen more commonly within missense than nonsense mutations. Missense mutations more frequently showed ASE in oncogenes and tumour suppressor genes than essential genes, whilst nonsense mutations showed ASE more frequently in tumour suppressor genes than essential genes (Figure 1E). In this context, repression of the wild-type allele of a canonical tumour-suppressor gene represents a copy-number independent mechanism of allelic inactivation, and highlights that ASE can generate functional variation in tumours^18^.

Thus, measurements of the clonal status of the mutation, allelic imbalance and allele-specific expression all exhibit differences between cancer genes and non-cancer genes, and provide rationale for the use of RNA VAF to identify cancer genes. Indeed, clonal mutations, mutations in allelic imbalance and mutations showing allele-specific expression exhibited elevated RNA VAF (Figure 1F).

We therefore tested the ability of RNA VAF to differentiate between cancer genes and non-cancer genes. We considered five canonical cancer genes and five non-cancer genes in colonic and rectal tumours from TCGA. Clear differences were observed comparing the distributions of RNA VAFs between cancer genes and non-cancer genes, with non-cancer genes tending to harbour lower RNA VAFs than cancer genes (Figures 1G).

We then assessed whether RNA VAF might encode differences between functional and non-functional mutations within putative cancer genes. To assess this signal, we considered mutations in CGC genes within colorectal cancer. We postulated that relative differences in RNA VAF between non-synonymous and synonymous mutations within cancer genes might represent a signal of selection captured by this metric. Indeed, there was a significant difference in RNA VAF between non-synonymous and synonymous within CGC genes (Figure 1H, Wilcoxon *p*<2.2e-16). This difference was more pronounced than when considering DNA VAF (Wilcoxon effect size 0.16 compared to 0.08; Figure S5).

### RVdriver

To extend this observation genome-wide and leverage RNA VAF information for the identification of cancer genes, we developed an approach, *RVdriver*, to test the observed RNA VAFs of mutations against an expected background (Methods) and implemented this across each of the cancer types within TCGA.

Briefly, within the cohort of tumours of each cancer type, we considered each gene surpassing a threshold of sufficient nonsynonymous mutations required to test against an expected background. This threshold was a minimum of 4 mutations within a given gene at a minimum required depth of 8 non-duplicated reads, across all tumours of a specific cancer types. A background set of mutations was generated for each comparison. We utilised a consistent number, 10, of synonymous mutations sampled from across a range of genes within each patient with a mutation within a gene of interest and tested their RNA VAFs against those of the nonsynonymous mutations within that gene. Sampling of consistent numbers of synonymous mutations across tumours was performed to minimise the potential confounding impact of inter-tumour differences in the distribution of RNA VAFs. Furthermore, the impact of lowly expressed genes was mitigated by filtering out synonymous mutations with an RNA depth < 8 non-duplicated reads (Methods).

A mixed effects linear model was then computed, comparing RNA VAFs for the set of nonsynonymous mutations within the gene of interest to those of the ‘background’ set of synonymous mutations from across genes, with the tumour in which the mutation was identified added as a random factor. This selection of synonymous mutations and statistical test was bootstrapped 50 times, in order to limit the impact of sampling given VAFs are variable across mutations, and the resulting p value adjusted for repeated measures across the different genes tested within the cancer type to obtain a q-value for each gene. Genes with a q-value<0.01 were considered to be putative cancer genes, whilst those with a q-value>0.01 but <0.05 were considered to be ‘low-confidence’ putative cancer genes.

### Performance of *RVdriver* relative to genomic approaches

We applied *RVdriver* to our cohort of 7,948 tumours. Across the 30 cancer types, it identified 136 instances of CGC genes (here, as elsewhere^7^ used as a proxy for the true-positive rate) acting as putative cancer genes in a cancer type, compared to 50 non-CGC genes, highlighting its ability to identify meaningful biological patterns.

We next sought to understand whether our approach provided additional information in the context of cancer gene identification compared to methods leveraging DNA data alone. In order to do this, we computed output from four tools, *dNdScv*^*5*^, *mutPanning*^*7*^, *oncodriveCLUSTL*^*9*^ and *oncodriveFML*^*14*^, which were run separately on the cohort of tumours for which paired DNA and RNAseq was available. These tools, respectively, consider the difference between synonymous and nonsynonymous mutations at their putative sites; the nucleotide context of mutations; clustering of mutations around putative hotspots; and the predicted functional impact of mutations. We evaluated the relative performance of these tools by assessing the number of CGC genes compared to non-CGC genes discovered by each tool (using thresholds specified by each tool, see Methods). Within our cohort, *RVdriver* exhibited comparable performance relative to other tools, with a pan-cancer area under the ROC curve (AUC) of 0.79 at a q-value threshold of 0.01 (Figure 2A, 2B, 2C, S6, S7). AUC at a q-value threshold of 0.05 was 0.71 (Figure S8C). Of note, given a minimum requirement of four mutations per cancer type, it is possible that our approach would be expected to discover more CGC genes than a random background, given the propensity of these genes to be more frequently mutated. To that end, we computed results for this analysis but only considering frequently mutated genes; when analysed in this way, *RVdriver* maintained comparable performance relative to other tools across the majority of cancer types, although was worse in bladder and endometrial tumours (Figure S9, S10).

**Figure 2.**
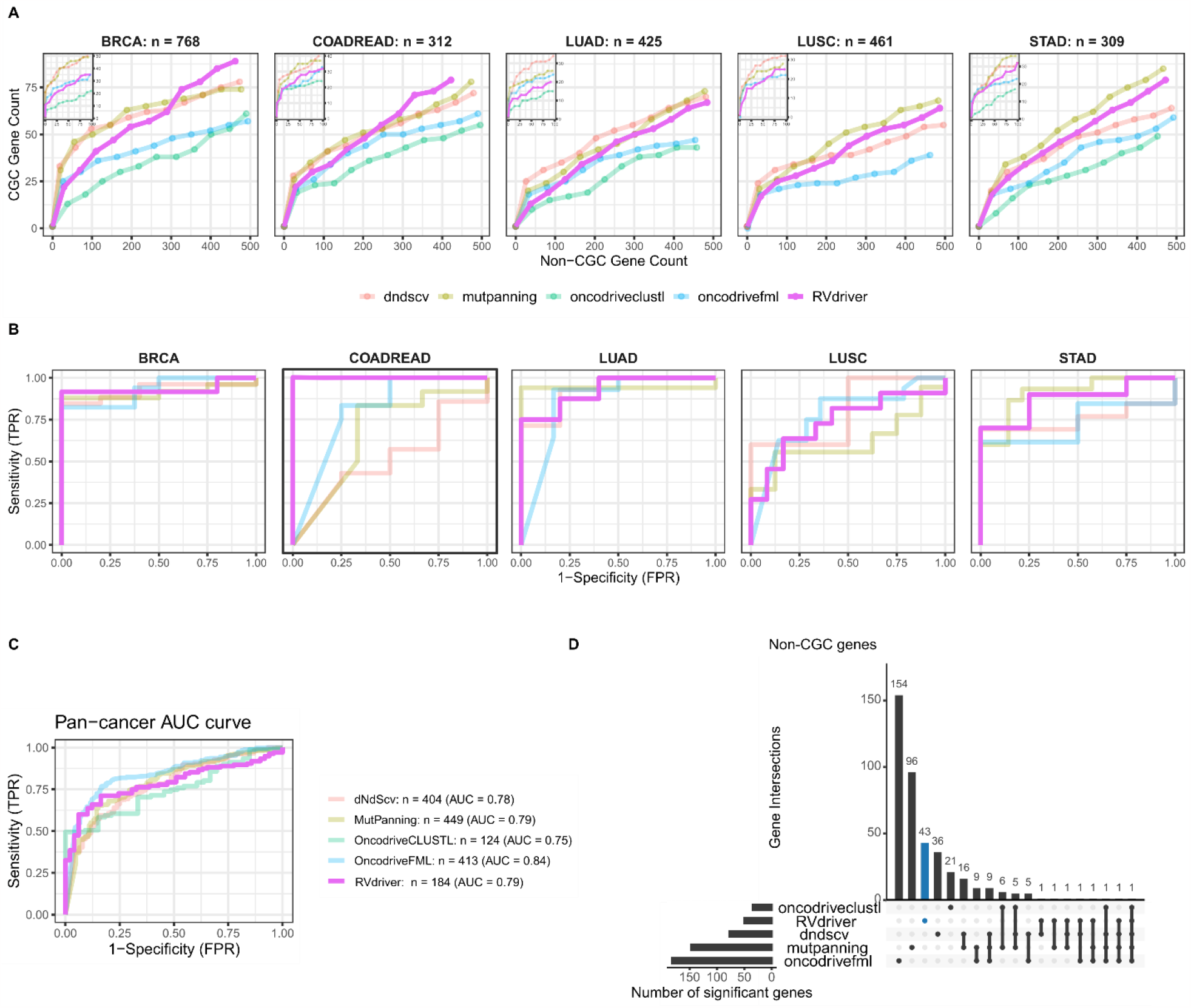
Performance of *RVdriver* identifying cancer genes. **A**. Benchmarking of *RVdriver* in five cancer types (BRCA = breast carcinoma; COADREAD = colorectal carcinoma; LUAD = lung adenocarcinoma; LUSC = lung squamous cell carcinoma; STAD = gastric adenocarcinoma) against four other established tools leveraging DNA information, *dNdScv, mutpanning, oncodriveclustl*, and *oncodriverfml*. Genes within the COSMIC cancer gene census list were treated as true positive results, and other genes as true negative results. The figure displays the number of CGC genes (y-axis), versus non-CGC genes (x-axis), identified within the top hits by the tools. **B**. ROC curves for CGC gene discovery for the same five cancer types. **C**. Area under the ROC curve for the identification of CGC genes across 30 cancer types, calculated independently for *RVdriver, dNdScv, mutpanning, oncodriveclustl*, and *oncodriverfml*. Number of genes identified as putative cancer genes according to tool-specific threshold annotated. **D**. UpSet plot showing overlap between the non-CGC genes discovered by *RVdriver* and the four DNA approaches. *RVdriver* here identifies forty three instances of non-CGC genes acting as cancer genes within a cancer type that were not highlighted by the four DNA approaches.

We studied the degree to which putative cancer genes identified by *RVdriver* overlapped with those identified by *dNdScv*^*5*^, *mutPanning*^*7*^, *oncodriveCLUSTL*^*9*^ and *oncodriveFML*^*14*^ (Figure 2D, S11). Considering only CGC genes (Figure S9), *RVdriver* was able to identify 14 separate gene-cancer type interactions that were not picked up by the four tools analysing only the DNA, as well as a further 122 gene-cancer type interactions that were identified by at least one other tool. Furthermore, within non-CGC genes (Figure 2D), which might plausibly include some hitherto-undetected *bona fide* cancer genes, *RVdriver* described 43 putative cancer gene-cancer type relationships which were not identified as such by other tools, as well as 7 putative cancer gene-cancer type relationships which had support from at least one other tool.

### Putative cancer genes identified by *RVdriver*

The output from *RVdriver* across the 30 cancer types in TCGA is displayed in Figure 3. This analysis highlights specific cancer genes that were identified by the MC3 group within a near-identical (excluding approximately 1,300 samples with DNA information available only) dataset^20^, as well as CGC genes.

**Figure 3.**
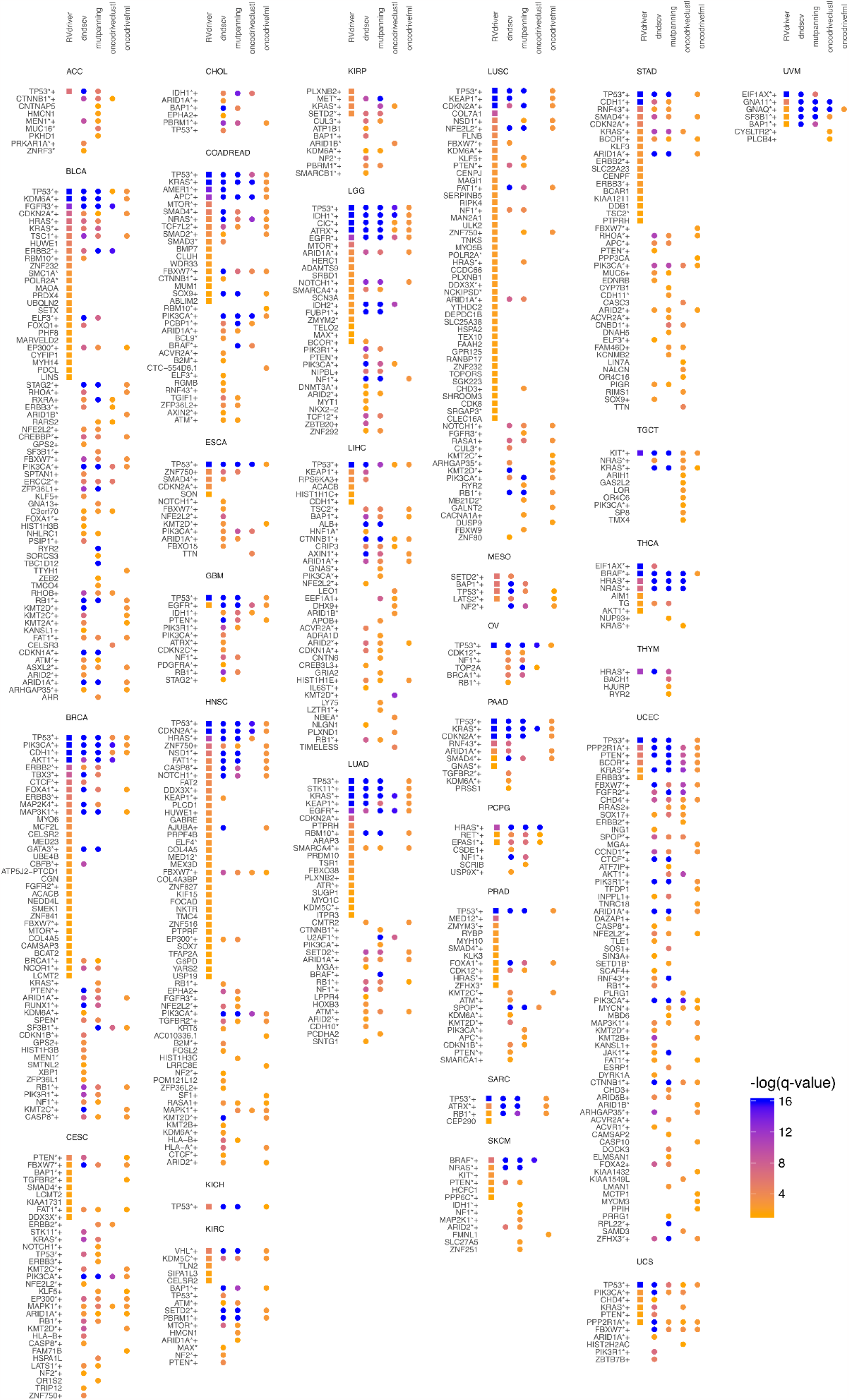
Cancer genes identified by *RVdriver* and DNA approaches. Using paired whole-exome and RNA sequencing from 7,948 tumours across 30 cancer types, *RVdriver, dNdScv, mutpanning, oncodriveclustl*, and *oncodriverfml* were used to identify putative cancer genes. Point colour indicates the negative log of the q value outputted by the tool. Only genes with a q value of less than 0.05 in one or more tool are plotted. **+** indicates gene identified as significant by Bailey et al within that cancer type. **\*** indicates gene contained within the CGC list.

Many canonical cancer genes were identified using *RVdriver*. For example, *TP53* is highlighted as a cancer gene in 19 cancer types. However, there are other putative cancer genes identified by *RVdriver* that are not identified by the DNA tools. These include cancer genes that were not highlighted as such within that cancer type, such as: *ERBB3, FGFR2* and *FBWX7* (breast); *SMAD4* and *BAP1* (cervical); *MTOR* (colorectal); *CDKN2A* (oesophageal); *DDX3X* (head and neck); *MTOR* and *MAX* (glioma); *CDH1* (hepatocellular carcinoma); *CDKN2A, KDM5C* and *ATR* (lung adenocarcinoma); *KDM6A* (lung squamous cell carcinoma); *MED12, SMAD4* and *HRAS* (prostate); *PPP6C* and *KIT* (melanoma); *CDKN2A* and *ERBB2* (stomach); *AKT1* (thyroid); and *ERBB3* (endometrial). Importantly, *RVdriver* also implicated several novel cancer genes that would be worthy of further investigation and analysis to determine their role in cancer development and progression.

### RNA VAF can predict driver and passenger mutations within established cancer genes

Given the ability of *RVdriver* to identify established and putative cancer genes, we investigated whether the underlying data, namely the RNA variant allele frequencies of somatic mutations, might enable functional driver mutations to be distinguished from non-functional passenger mutations in cancer genes. Distinguishing between driver and passenger mutations has been a significant research focus, and efforts such as Intogen have utilised machine learning approaches leveraging multiple features that describe tumourigenesis within different genes and cancer types. This has resulted in detailed atlases of probable driver mutations within putative cancer genes, including saturation analyses generated from next generation DNA sequencing data that outperform experimental mutation saturation analyses, notably BoostDM^15^.

We first sought to establish the degree to which RNA VAF was able to independently identify putative driver and passenger mutations as defined by BoostDM. RNA VAF was computed for each mutation with a minimum depth of 8 non-duplicated reads at that position within individual cancer types. Within missense mutations in tumour suppressor genes, there was a strong significant correlation between the driver/passenger annotation in BoostDM and the RNA VAF of the mutation, whilst a weak yet significant correlation was seen within such mutations in oncogenes (Figure 4A, 4B). Similar relationships were not observed in nonsense mutations, in keeping with the tiny fraction of such mutations that represent probable passengers in tumour suppressor genes, or drivers in oncogenes.

**Figure 4.**
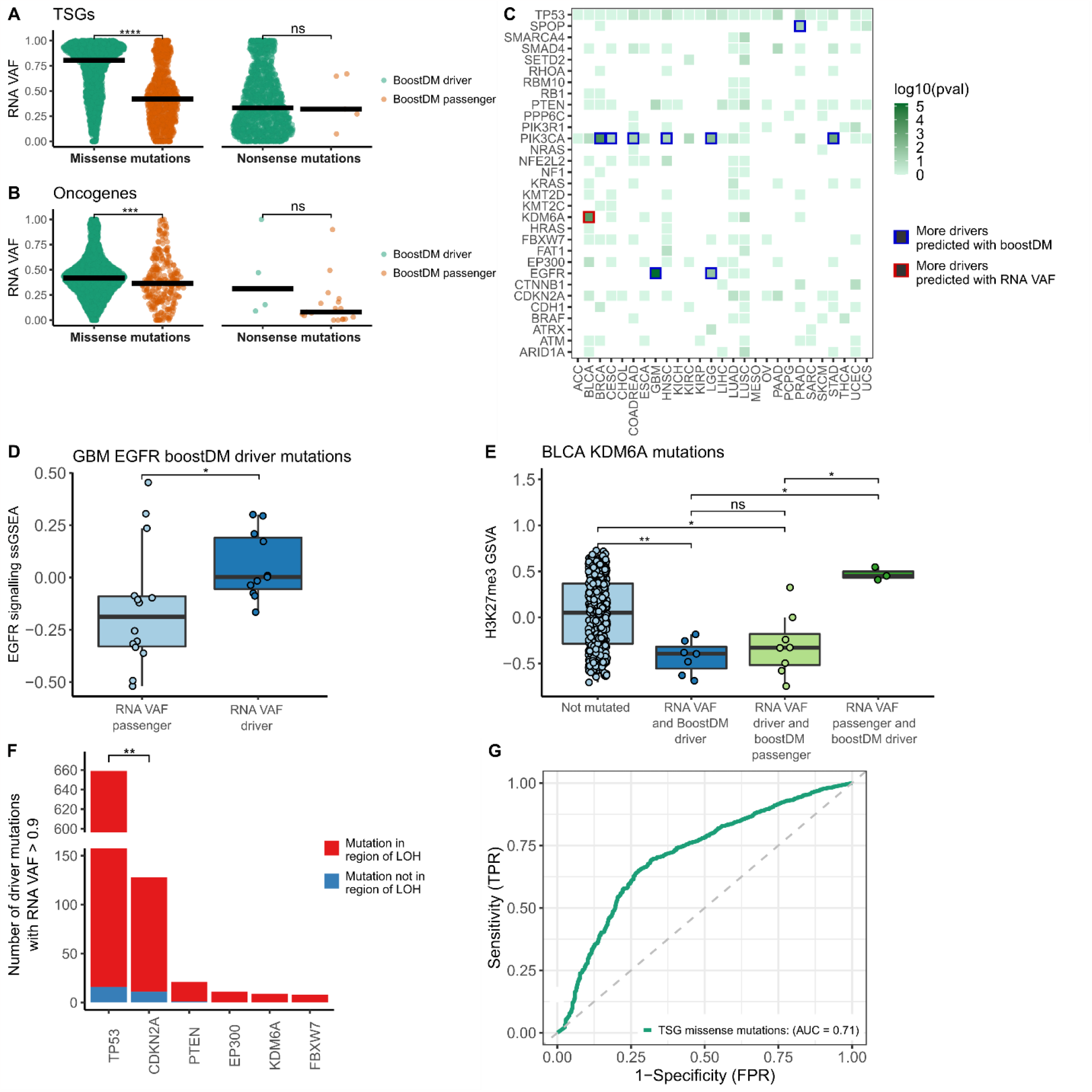
RNA VAF in driver and passenger mutations within established cancer genes. **A**. RNA VAFs of mutations identified within a mutation saturation analysis of cancer genes^15^. Cancer genes were split by oncogene/tumour suppressor gene status, and whether they were missense or nonsense mutations. Mutations are coloured by BoostDM driver status (green: BoostDM driver; orange: BoostDM passenger). **B**. Focussed analysis of concordance of classification of driver and passenger mutations between BoostDM and RNA VAF. Genes harbouring more (p<0.1) mutations identified as drivers by BoostDM, and passengers by RNA VAF (i.e. having an RNA VAF < 0.1) have a blue outline. Tile colour indicates the negative log of the p value for significance of Fisher’s exact test, comparing the number of driver mutations and passenger mutations identified within the gene by BoostDM and RNA VAF. **C**. ssGSEA scores for REACTOME ‘EGFR signalling in cancer’ gene set among glioblastomas harbouring a BoostDM driver, split by RNA VAF < 0.1 or RNA VAF > 0.9. Wilcoxon test was used to estimate significance. **D**. ssGSEA scores for the ‘altered tri-methylation of lysine 27 on histone H3’ gene set from Schlesinger et al^.38^ among bladder carcinomas. These are split by whether they harbour a *KDM6A* driver mutation, as defined by BoostDM or RNA VAF (passenger: RNA VAF < 0.1 or driver: RNA VAF > 0.9). Wilcoxon test was used to estimate significance. **E**. High RNA VAF (>0.9) mutations within commonly mutated tumour suppressor genes. The majority of such mutations were found within a region of loss-of-heterozygosity (LOH). However, a small fraction of such instances are LOH-independent. More such instances were observed within *CDKN2A* than *TP53* (Fisher’s exact test). **F**. RNA VAFs were computed for mutations identified within the BoostDM *in silico* saturation analysis of cancer genes. 8,716 mutations were expressed with more than 8 reads at that position. Scores from BoostDM were utilised as ground truth. Thus, an AUC of 1 would indicate perfect alignment between RNA VAF and BoostDM scores.

Given that there are multiple competing factors that influence the RNA VAF of a somatic mutation, it is possible that further information of biological interest might be encoded within cases where there is disagreement between BoostDM and RNA VAF predicted driver mutations. We therefore sought to evaluate such cases. Among all mutations, 308 were predicted to be drivers by BoostDM, and high-confidence passengers by RNA VAF (i.e. RNA VAF less than 0.1), whilst the reverse was true (BoostDM passengers with RNA VAF greater than 0.9) in 117 cases.

We performed a Fisher’s exact test for each cancer type to compare the numbers of putative driver and passenger mutations within each gene and cancer type identified by BoostDM and RNA VAF. Results for each gene and cancer type combination are displayed in Figure 4C, where genes and cancer types for which BoostDM and RNA VAF disagreed significantly about the numbers of driver and passenger mutations are outlined in blue or red.

There were nine instances where the RNA VAF classified significant numbers of mutations as passengers where BoostDM had classified them as drivers. All of these were in oncogenes. For example, of fifty five BoostDM driver mutations within *EGFR* in glioblastoma multiforme, nineteen had RNA VAF < 0.1, compared to only eleven with RNA VAF > 0.9. This either suggests that these mutations are not *bona fide* drivers, or that within cancers harbouring such a mutation, there is an absence of selective pressure for elevated expression of the mutated allele.

Within tumours containing a BoostDM *EGFR* driver mutation and an RNA VAF > 0.9, ssGSEA (single sample gene set enrichment analysis) revealed significantly higher expression of the REACTOME ‘EGFR signalling in cancer’ gene set, when compared to BoostDM driver mutations with an RNA VAF < 0.1 (*P*=0.025, Figure 4C).

Furthermore, of the thirty tumours with BoostDM *EGFR* driver mutations, where RNA VAF was either greater than 0.9 or less than 0.1 (i.e. a confirmed positive or negative call), twenty-seven (90%) harboured ultra-high copy-number amplifications (total copy-number > 8; described in the literature as consistent with the presence of ecDNA^35,36^). This highlights that in the context of this mutation, there is not a consistent selection pressure for amplification of the mutation.

Similarly, six of the nine instances of RNA VAF repeatedly classifying BoostDM drivers as passenger mutations involved the gene *PIK3CA* (i.e. this happened separately across six cancer types). This either suggests that *PIK3CA* mutations are not *bona fide* drivers or adds to evidence supporting its activity acting in a dose-dependent manner, with cancers only being able to tolerate a certain level of mutated *PIK3CA* activity^37^.

There was one instance (i.e. a combination of gene and cancer type) where RNA VAF repeatedly identified (Fisher’s exact test, p<0.1) putative driver mutations among BoostDM passengers: *KDM6A* in urothelial carcinoma of the bladder. Here, it is plausible that BoostDM might have underestimated the putative functional impact of these mutations. In order to test this hypothesis, we computed ssGSEA scores for the impact of *KDM6A* across bladder tumours: altered tri-methylation of lysine 27 on histone H3 (H3K27me3)^38^. This analysis revealed a consistent trend towards reduced H3K27me3 gene set activity in tumours with mutations that BoostDM and RNA VAF agreed were drivers (i.e. RNA VAF > 0.9), relative to those without *KDM6A* mutations (Figure 4D). However, where RNA VAF disagreed with BoostDM, gene set activity suggested that RNA VAF results tended to be more consistent with tumour phenotype than BoostDM (i.e. in BoostDM passengers that were RNA VAF drivers, H3K27me3 activity was similar to BoostDM drivers that were RNA VAF drivers; whilst in BoostDM drivers that were RNA VAF passengers, H3K27me3 activity was more similar to non-*KDM6A*-mutated tumours).

Significantly elevated RNA VAF (i.e. >0.9) in tumour suppressor genes is consistent with non-expression of the wild-type allele. Canonically, this can result from loss-of-heterozygosity (LOH) of the wild-type allele. However, allele-specific copy-number profiling from SNP array data allows identification of instances where wild-type allele inactivation occurs independently of LOH (i.e. its expression is repressed). The frequency of LOH-independent inactivation of the wild-type allele was compared across tumour suppressor genes. Whilst for many genes there were not sufficient available mutations to power this analysis, this did reveal that LOH-independent wild-type allelic inactivation was more common in *CDKN2A* than *TP53* (Figure 4F).

Finally, we evaluated the performance of RNA VAF as a metric for identifying single driver mutations within cancer genes by comparing its sensitivity and specificity to scores from BoostDM from Intogen as a set of true positive calls.

We considered all (8,716) mutations where RNA coverage was 8 or more reads within the 68 genes analysed by BoostDM, using the median RNA VAF of mutations that recurred within cancer types. The AUC for all mutations was 0.59. However, this was not evenly distributed across all mutation types. For example, the vast majority of nonsense mutations within tumour suppressor genes were considered by BoostDM to be driver mutations. However, for missense mutations within tumour suppressor genes, where functional impact is typically less certain, the AUC of RNA VAF was 0.71. This indicates that RNA VAF encodes significant information about the functional impact of mutations within cancer genes. Of note, within oncogenes the AUC for missense mutations was 0.53, and for nonsense mutations it was 0.78. When restricting this analysis to mutations with higher confidence of driver/passenger status from RNA VAF (i.e. with RNA VAF < 0.1 or RNA VAF > 0.9), the ability to predict BoostDM scores improved. Among such missense mutations in oncogenes, the AUC was 0.61; and for missense mutations in tumour suppressors it was 0.75. This highlights that RNA VAF might be less useful in predicting putative driver mutations within oncogenes than tumour suppressor genes.

## Discussion

The identification of cancer genes has been an important exercise, leveraging patterns of mutagenesis within tumour genomes to identify signals of positive selection and recurrence within cancer genes. However, to date, the role for bulk RNA sequencing in this process has been limited.

Here, we have proposed a novel approach for the identification of cancer genes. *RVdriver*, which utilises RNA variant allele frequencies from somatic mutations, identifies clear signals in patterns of mutation expression between cancer genes and non-cancer genes. It exhibits comparable performance to DNA-based tools, and importantly identifies putative novel cancer genes, not highlighted to date by DNA approaches, that might merit further investigation.

There seem to be certain cancer genes which appear to be more easily discovered by *RVdriver* than DNA-based tools. For example, *MTOR*, which is an established cancer gene, is identified as such by *RVdriver* in low-grade glioma, colorectal carcinoma, and breast carcinoma, but not by DNA-based tools in any of those instances. Indeed, *RVdriver* highlights 30 instances of genes within the COSMIC cancer gene census list acting as cancer genes within cancer types which are not picked up by DNA approaches. Similarly, *RVdriver* identifies 118 instances of genes outside of the COSMIC cancer gene census list acting as cancer genes. It is possible that some such genes, particularly those called with low confidence (0.01<q<0.05), might represent false positives. However, other candidates, including *COL7A1*, a component of collagen in lung squamous cell carcinoma; *CLUH*, a regulator of mitochondrial metabolism in colorectal carcinoma and lung squamous cell carcinoma; and *CENPF, CEP290* and *CENPJ*, encoding centrosomal proteins in gastric carcinoma, sarcoma and lung squamous cell carcinoma respectively, might well be worthy of further investigation.

Conversely, *RVdriver* fails to identify a number of established cancer genes in certain instances. Whilst this clearly, and perhaps unsurprisingly, indicates that *RVdriver* is not a panacea in terms of its ability to identify cancer genes, it might also provide insight into important tumour biology. For example, *RVdriver* fails to identify *PIK3CA* as a putative cancer gene in thirteen tumour types in which it is identified by DNA approaches, and only successfully identifies it as such in two cancer types. The possibility that *PIK3CA* might represent an outlier in terms of its patterns of mutation expression is supported by orthogonal analysis, in which *PIK3CA* mutations across six cancer types exhibit low RNA VAF despite being predicted by BoostDM to be driver mutations. This highlights that *RVdriver* can reveal facets of tumour biology that might be missed by existing approaches.

A clear limitation of *RVdriver* is that it is limited by necessity to discovery within frequently mutated genes. Such genes are already enriched for cancer genes. Larger datasets would provide power to assess the functionality of less frequently mutated genes also.

We also demonstrate the RNA VAF of mutations can highlight differences between driver and passenger mutations among missense mutations within tumour suppressor genes. It could therefore act as an important orthogonal adjunct to attempts seeking to distinguish between functional and non-functional mutations in cancer. Indeed, by phenotyping tumours according to relevant gene set expression scores, we show that RNA VAF identifies instances of possible driver-passenger misclassification events within the cancer genes *EGFR* and *KDM6A*.

RNA VAF likely reflects the activity of different processes in tumours. First, somatic mutations that are more likely to be associated with a clonal sweep (i.e. driver mutations) are more likely to be present in a greater fraction of cancer cells, and hence exhibit higher RNA VAF. Second, in many oncogenes, amplification of the mutated allele is associated with increased oncogenic activity; whilst retention of a mutated allele in the context of a loss-of-heterozygosity is classical in tumour suppressor genes. Third, in tumours, expression preferentially is derived from the tumour component of the admixed sample^26^, and particularly so for many cancer genes which are often highly expressed in epithelial tissues. Fourth, non-mutational mechanisms underpinning allele-specific expression, such as methylation, might plausibly favour the non-mutated allele in the presence of a driver mutation^31^.

Irrespective of the mechanism underpinning this phenomenon, we have presented a simple approach, encapsulated in *RVdriver*, that adds value to analyses seeking to identify new cancer genes. This may be of value to genomic-transcriptomic analyses of cancer genes; and future studies should also attempt to interrogate the genome and transcriptome in parallel to understand in full the landscape of functional variation in tumours.

## Methods

### Download of TCGA data

RNA sequencing data (BAM files and processed gene expression counts) for 7,948 samples across 30 cancer types was downloaded via the GDC data portal (https://portal.gdc.cancer.gov/) using the GDC data transfer tool. For a full list of samples and the specific RNA-seq bam file used in this analysis, see Table S1.

### Mutation data

A publicly available mutation annotation file (MAF) file compiled by the MC3 working group^21^ was downloaded (https://gdc.cancer.gov/about-data/publications/mc3-2017). This MAF file was filtered to include only the samples investigated in this study (Table S1). In addition, variants assigned as ‘PASS’ were selected, removing potentially artifactual variants. In Ovarian cancer (OV), variants that had only the ‘wga’ filter were also kept as samples within this cancer type were primarily sequenced using whole genome amplification. This is in keeping with previous work^20^. Samples that had been excluded from previous studies based on pathology review, samples excluded from previous studies as ‘hypermutators’ or samples flagged as having degraded RNA were also removed^20^. Finally, mutations lying within the ENCODE blacklisted regions were also removed^22^.

### Determining the RNA VAF of somatic mutations

Each BAM file was first pre-processed following GATK’s best practices, marking and removing duplicate reads and applying base quality score recalibration (GATK v4.1.7.0) using default parameters. After lifting over the position (using the liftOver R package) of each mutation from hg19 to hg38, the RNA reference and alternate counts for each position containing a single nucleotide variant were calculated using the bam2r function from the R package deepSNV using default parameters (v1.42.0)^39^. The RNA variant allele frequency was calculated as the sum of non-duplicated reads aligning to the alternate allele divided by the sum of non-duplicated reads aligning to both the reference and alternate alleles at that position.

### RVdriver

RVdriver was run separately on individual cancer types and involved the following steps.

1. Filter the MAF file for synonymous and nonsynonymous single nucleotide variants only (non-synonymous mutations comprised nonsense, missense, splice site and non-stop mutations).
2. Calculate which genes to test within a given cancer type. A gene was tested if, within a single cancer type, it had > 3 non-synonymous mutations at an RNA depth > 7 non-duplicated reads.
3. Iterate through each of these genes, performing the following test.
  a. Filter for all non-synonymous mutations within the gene of interest
  b. Sample a consistent number (10) of synonymous mutations (at an RNA depth > 7 non-duplicated reads) from each tumour sample containing a nonsynonymous mutation within the gene of interest.
  c. Generate a mixed effects linear model comparing the RNA VAF distribution of nonsynonymous mutations within the gene of interest against the ‘background’ set of synonymous mutations using the lmerTest R package and the following formula:

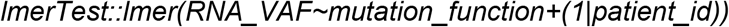
  d. By using patient_id as a random effect, we control for differences in sample-specific variables such as estimated tumour content as well as differences in overall expression from the tumour compartment relative to the non-tumour compartment of the admixed bulk sample.
  e. Bootstrap steps b and c 50 times to control for the high variability of RNA VAFs sampled across and within patients.
  f. Take the median T value and degrees of freedom from the bootstrapping approach. Use the following function to derive a p value.

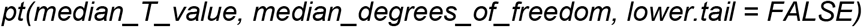
  g. Perform multiple hypothesis testing correction on all gene results using the Benjamini-Hochberg approach. A gene is classified as a putative driver with high confidence if the q value is < 0.01 and lower confidence if the q value is 0.01 < q < 0.05.

Code to run RVdriver is available here: https://github.com/McGranahanLab/RVdriver

### Classification of known cancer genes

Cancer genes were classified as those present in the Cancer Gene Census (CGC) list (v.96)^23^. This classification was used in the analyses contained within figures 1, 2 and 3.

Oncogenes and Tumour suppressor genes (TSGs) were assigned as outlined in a previous pan cancer analysis^20^. In cases where a gene had a cancer specific oncogenic or tumour suppressive function, this was prioritised over the pan cancer assignment for that gene^20^. This classification was used within the analyses conducted within figures 1,3 and 4.

### Somatic copy number assignment

Affymetrix SNP 6.0 arrays were downloaded from the TCGA (see above). ASCAT was run to obtain allele specific copy numbers as well as purity and ploidy estimates for each sample^34^.

### Estimating the cancer cell fraction of mutations

The cancer cell fraction (CCF) was estimated using the following formula as per McGranahan et al^29^.

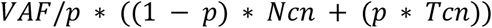

Where *p* is the tumour purity derived using ASCAT, *Ncn* is the normal copy number at the mutated position (assumed to be 2 within this analysis) and *Tcn* is the copy number within the tumour at the mutated position, derived from ASCAT. Point estimates for CCF and their associated confidence intervals were then calculated using a binomial distribution as previously described^29^. A mutation was defined as clonal when the 95% confidence interval overlapped 1, and subclonal when this was not the case.

### Estimating the mutant copy number fraction

The mutant copy number (the number of chromosomal alleles harbouring the mutation) was estimated using the following formula.

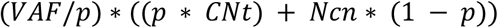

The mutant copy number fraction was estimated by dividing the mutation copy number by the total copy number within the tumour at the mutated position, and capped at 1.

### Identifying allele specific expression

Allele specific expression was calculated using the following approach. In accordance with the expected distribution of allelic expression, a beta-binomial test was used to test for allele-specific expression^40^ using the pbetabinom function from the R package VGAM (v1.1.1)^41^ and an over-dispersion parameter of 0.05. To assess the probability of observing allele specific read counts at least as disparate as the observed distribution given the estimated mutant copy number and the total copy number, the following beta binomial test was performed.

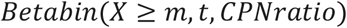

Where m represents the RNA-seq read counts of the mutated allele, t is the total RNA reads at that position and CPNratio is the estimated mutant copy number divided by the total copy number at the mutated position. Genes with a p value < 0.05 from this test were considered to show copy number independent allele-specific expression.

### DNA based tools to detect cancer genes

Four different tools utilising patterns observed within genomic data were used to benchmark the performance of RVdriver. All 4 tools were run on default settings and on all mutations available for a given sample.

1. dNdScv looks for excess numbers of nonsynonymous mutations at nonsynonymous sites compared to synonymous mutations at synonymous sites to infer positive selection for mutations in putative cancer genes^5^.
2. OncodriveFML uses scores for the deleteriousness of mutations (in this case CADD scores^42^) to identify genes where there are an excess of more functional mutations in a gene than would be expected by chance^14^.
3. OncodriveCLUSTL looks for clustering of mutations in specific hotspots within a given gene, identifying putative drivers as those that have an increased number of hotspots than would be expected by chance^9^.
4. MutPanning identifies cancer genes as those with an excess number of mutations above the background rate as well as an excess number of mutations in unusual trinucleotide contexts^7^.

When benchmarking RVdriver against these tools, we used q value thresholds to call putative driver genes as outlined in each respective publication. These were as follows:

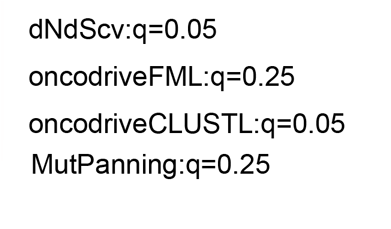

After calling putative drivers across these thresholds, area under the ROC curves were generated for each tool using the R package ROCit (v2.1.1), using genes within the CGC-list as true positives.

We used the R package UpSetR^43^ (v1.4.0) to investigate the overlap between putative cancer-type-specific cancer genes determined by each tool at their respective q values. We split this analysis to investigate the overlap between each tool within cancer genes as defined by the CGC list and non-CGC list genes.

### Driver annotation using BoostDM

Boostdm is a machine learning method for in silico saturation mutagenesis. Utilising different features of mutations it has generated a cancer-type specific atlas of probable driver and passenger mutations^15^. This pan-cancer atlas was downloaded (https://www.intogen.org/boostdm/downloads) and wrangled to match tumour types with those present within our dataset. This resulted in a mutation table of 10,497 SNVs with matched, tumour type specific, boostDM driver/passenger annotation (binarised about a boostDM score of 0.5).

### Gene set enrichment analysis

To perform geneset enrichment analysis, BLCA and GBM expression count tables were downloaded from the TCGA portal (see above). Lowly expressed genes (where > 20% of the cohort had < 5 counts) were removed. This filtered counts table was then transformed into VST counts using the vst command from the R package DESeq2^44^ (v1.34.0).

To investigate the activity of EGFR signaling in GBM and KDM6A in BLCA, the reactome gene sets “REACTOME_SIGNALIN_BY_EGFR_IN_CANCER” and “SCHLESINGER_H3K27ME3_IN_NORMAL_AND_METHYLATED_IN_CANCER”^38^ were downloaded respectively from the reactome pathway knowledgebase^45^. We performed a single sample GSEA (ssGSEA) for the above gene sets using the R package fgsea (v1.10.1)^46^ on VST counts using a Gaussian distribution and with default parameters.

### Analyses

RVdriver was run in the R environment (v3.5.2). To account for differences in sample-specific variables within RVdriver (see above), we use linear mixed effects models fitted using the package lmerTest (v3.1.0).

All plots were generated in the R environment (v4.2.0) using ggplot2 (v3.3.6), cowplot (v1.1.1), ggpubr (v0.4.0) and ggbreak (v0.1.1). All Wilcoxon tests performed are two-sided, using the function wilcox.test() in base R, with the effect size obtained using wilcox_effsize() from the package rstatix (v0.7.0).

## Supplementary figure legends

Supplementary figure 1: Cohort overview. Bars represent the number of samples with paired DNA/RNA-sequencing data across the available cancer types within the TCGA cohort.

Supplementary figure 2: A. Proportion of expressed mutations across all mutations within the cohort mutation table. A mutation was considered expressed when the alternate allele had 2 or more non-duplicated reads aligned at the mutated position. A mutation was considered non-expressed when the total RNA depth at the position was >= 2 non-duplicated reads and the RNA alternate read count was < 2 non-duplicated reads. The gene in which the mutation was found was considered not expressed when the RNA depth at the mutated position was < 2 non-duplicated reads. B. Portion of expressed mutation across different mutation types and across genes present or not present in the Cancer Gene Census list. Definitions for mutant expression are the same as above.

Supplementary figure 3: Proportion of expressed mutations within genes in the COSMIC cancer gene census list, and across different mutation types and cancer types. Definitions for mutant expression are the same as figure S2.

Supplementary figure 4: Proportion of expressed mutations within genes not in the COSMIC cancer gene census list, and across different mutation types and cancer types. Definitions for mutant expression are the same as figure S2.

Supplementary figure 5: DNA VAF of synonymous versus non-synonymous mutations within COSMIC cancer gene census genes; **** indicates p<0.0001; Wilcoxon test.

Supplementary figure 6: Benchmarking of *RVdriver* across all cancer types against four other established tools leveraging DNA information, *dNdScv, mutpanning, oncodriveclustl*, and *oncodriverfml*. Genes within the COSMIC cancer gene census list were treated as true positive results, and other genes as true negative results. The figure displays the number of CGC genes (y-axis), versus non-CGC genes (x-axis), identified within the top hits by the tools until 500 non-CGC genes were reached.

Supplementary figure 7: Benchmarking of *RVdriver* across all cancer types against four other established tools leveraging DNA information, *dNdScv, mutpanning, oncodriveclustl*, and *oncodriverfml*. Genes within the COSMIC cancer gene census list were treated as true positive results, and other genes as true negative results. The figure displays the number of CGC genes (y-axis), versus non-CGC genes (x-axis), identified within the top hits by the tools until 100 non-CGC genes were reached.

Supplementary figure 8. A. Benchmarking of *RVdriver* in five cancer types (BRCA = breast carcinoma; COADREAD = colorectal carcinoma; LUAD = lung adenocarcinoma; LUSC = lung squamous cell carcinoma; STAD = gastric adenocarcinoma) against four other established tools leveraging DNA information, *dNdScv, mutpanning, oncodriveclustl*, and *oncodriverfml*. Genes within the COSMIC cancer gene census list were treated as true positive results, and other genes as true negative results. The figure displays the number of CGC genes (y-axis), versus non-CGC genes (x-axis), identified within the top hits by the tools. B. ROC curves for CGC gene discovery for the same five cancer types using a q value cut-off of 0.05 for RVdriver. C. Area under the ROC curve for the identification of CGC genes across 30 cancer types, calculated independently for *RVdriver, dNdScv, mutpanning, oncodriveclustl*, and *oncodriverfml* using a q value cut-off of 0.05 for RVdriver. n = Number of genes identified as putative cancer genes according to tool-specific threshold annotated. D. UpSet plot showing overlap between the non-CGC genes discovered by *RVdriver* and the four DNA approaches. *Rvdriver* here identifies 114 instances of non-CGC genes acting as cancer genes within a cancer type that were not highlighted by the four DNA approaches.

Supplementary figure 9: Benchmarking of *RVdriver* across all cancer types against four other established tools leveraging DNA information, *dNdScv, mutpanning, oncodriveclustl*, and *oncodriverfml*. Here, only genes tested across all tools are considered (i.e. a minimum of 4 mutations at an RNA depth > 7 non-duplicated reads within that gene and cancer type were required). Genes within the COSMIC cancer gene census list were treated as true positive results, and other genes as true negative results. The figure displays the number of CGC genes (y-axis), versus non-CGC genes (x-axis), identified within the top hits by the tools until 500 non-CGC genes were reached.

Supplementary figure 10: Benchmarking of *RVdriver* across all cancer types against four other established tools leveraging DNA information, *dNdScv, mutpanning, oncodriveclustl*, and *oncodriverfml*. Here, only genes tested across all tools are considered (i.e. a minimum of 4 mutations at an RNA depth > 7 non-duplicated reads within that gene and cancer type were required). Genes within the COSMIC cancer gene census list were treated as true positive results, and other genes as true negative results. The figure displays the number of CGC genes (y-axis), versus non-CGC genes (x-axis), identified within the top hits by the tools until 100 non-CGC genes were reached.

Supplementary figure 11: A. UpSet plot showing overlap between the CGC genes discovered by *RVdriver* and the four DNA approaches (using a q value of 0.01 for RVdriver). *Rvdriver* here identifies 14 instances of CGC genes acting as cancer genes within a cancer type that were not highlighted by the four DNA approaches. B. UpSet plot showing overlap between the CGC genes discovered by *RVdriver* and the four DNA approaches (using a q value of 0.05 for RVdriver). *Rvdriver* here identifies 33 instances of CGC genes acting as cancer genes within a cancer type that were not highlighted by the four DNA approaches.

## Acknowledgements

The results published in this work are based on data generated by the TCGA project established by the National Cancer Institute and National Human Genome Research Institute. The data were retrieved through database of Genotypes and Phenotypes (dbGaP) authorization (accession number phs000178.v11.p8). Information about TCGA and the investigators and institutions can be found at http://cancergenome.nih.gov/.

Figure 1B was created with BioRender.com

